# Metabolic Cost Asymmetry as a Structural Regulariser for Recurrent Spiking Neural Networks

**DOI:** 10.64898/2026.10.09.758141

**Authors:** Roberto C. Sotero, Jose M. Sanchez-Bornot

## Abstract

Cortical energy expenditure is heavily dominated by synaptic transmission, with excitatory (glutamatergic) signalling demanding substantially more ATP than inhibitory (GABAergic) signalling. Yet existing trainable spiking neural networks treat energy as a generic, synapse-type-agnostic penalty. Here, we embed this fundamental glutamate/GABA cost asymmetry as a Lagrangian penalty within a surrogate-gradient learning framework, and use it to train a Dale’s-law-compliant recurrent microcircuit on the Spiking Heidelberg Digits (SHD) benchmark. Because glutamatergic synapses are metabolically more expensive, a single scalar penalty applied uniformly to all weights automatically exerts stronger gradient pressure on excitatory pathways. We demonstrate that this differential pressure alone is sufficient to produce three major effects. First, it moves the network out of the suboptimal, inhibition-dominated regime of the unpenalised network, simultaneously improving accuracy and reducing total energy several-fold. Second, it induces an emergent pruning hierarchy in which excitatory connections are eliminated first, while inhibitory connections exhibit graded resistance. Third, it actively expands the dimensionality of the population code at moderate penalty strengths. These structural motifs remain robust across a two-dimensional sweep of five cost ratios and fifteen structural densities (75 distinct conditions). Ultimately, our results demonstrate that a single biophysical cost asymmetry, operating without auxiliary regularisers or architectural priors, is sufficient to organise sparse, pathway-specific circuit structure in recurrent spiking networks.

**Highlights:**

- Asymmetric metabolic costs (glutamate > GABA) act as structural regularisers in SNNs.
- Metabolic pressure escapes inhibition-dominated regimes, raising task accuracy.
- Cost asymmetry induces a strict, emergent pathway-specific pruning hierarchy.
- Moderate metabolic pressure expands the dimensionality of the population code.
- Circuit motifs remain robust across diverse structural densities and cost ratios.

## 1. Introduction

Learning in biological neural systems is fundamentally resource-constrained. The brain consumes roughly 20% of resting metabolic energy despite comprising only 2% of body mass, and synaptic signalling dominates this massive cost (Attwell and Laughlin, 2001; Harris, Jolivet and Attwell, 2012). Crucially, this expenditure is not uniform across synapse types. Excitatory (glutamatergic) transmission, encompassing vesicle cycling and postsynaptic depolarisation, accounts for the vast majority of signalling energy, while inhibitory (GABAergic) transmission is substantially cheaper (Sengupta, Stemmler, Laughlin and Niven, 2010; Sterling and Laughlin, 2015). At the single-synapse level, GABA uptake into astrocytes, unlike glutamate uptake, does not elicit a measurable metabolic response, providing direct experimental evidence for a fundamental bioenergetic asymmetry (Chatton, Pellerin, Bhatt, Bhatt and Magistretti, 2003). Updated cortical energy budgets confirm this hierarchy, with postsynaptic glutamate receptor activation alone accounting for roughly half of neocortical signalling energy (Howarth, Gleeson and Attwell, 2012).

These observations have motivated a long line of normative theories proposing that neural codes are fundamentally shaped by energetic constraints. For instance, Laughlin et al. (1998) showed that blowfly photoreceptor signalling operates near the physical limits of energy-efficient information transfer. Levy and Baxter (1996) derived conditions under which sparse, low-rate firing maximises information per unit energy, and Harris et al. (2012) extended this logic to cortical populations, arguing that synaptic efficacy and firing statistics are tuned to maximise bits per ATP molecule. Niven et al. (2007) confirmed experimentally that synapses with lower transmission capacities encode information more economically, while Sengupta et al. (2010) showed that action potential waveforms themselves appear mathematically optimised to minimise metabolic cost. In parallel, theoretical work has established that excitatory/inhibitory (E/I) balance promotes metabolic efficiency by producing fewer, more informative spikes (Sengupta, Laughlin and Niven, 2013), and that this balance constrains population responses to low-dimensional manifolds whose geometry closely reflects task demands (van Vreeswijk and Sompolinsky, 1996; Denève and Machens, 2016; Langdon and Engel, 2023; Litwin-Kumar and Doiron, 2012). Despite this extensive body of work, the consequences of embedding synapse-type-specific ATP costs into a trainable learning objective for spiking networks remain unexplored.

Modern surrogate gradient methods (Neftci, Mostafa and Zenke, 2019; Zenke and Vogels, 2021) have enabled the end-to-end optimisation of recurrent spiking neural networks (SNNs) via backpropagation through time (BPTT), effectively integrating deep-learning methodologies into the spiking domain. Bellec et al. (2018) demonstrated that sparsely connected spiking networks can match Long Short-Term Memory (LSTM) networks performance on temporal tasks, and their subsequent e-prop rule (Bellec, Scherr, Subramoney, Hajek, Salaj, Legenstein and Maass, 2020) showed that biologically plausible approximations to BPTT can succeed in recurrent architectures. Furthermore, Cramer et al. (2022) introduced the Spiking Heidelberg Digits (SHD) benchmark, which converts spoken digits into multi-channel spike trains via a biologically inspired artificial cochlea model, and demonstrated surrogate gradient learning on neuromorphic systems. Eshraghian et al. (2023) subsequently consolidated these advances into the `snntorch` framework. On the efficiency side, Stanojevic et al. (2024) demonstrated high-performance deep spiking networks operating at merely 0.3 spikes per neuron per input, and Rathi and Roy (2023) introduced learnable membrane parameters that jointly optimise accuracy and spike count. However, all of these approaches employ generic regularisers (such as spike-count penalties, firing-rate targets, or *L*_1_ weight decay) that treat every synapse identically, entirely bypassing the biophysical mechanisms that make glutamatergic synapses fundamentally more expensive than GABAergic ones. Recent work has begun to explore energy-dependent plasticity rules (Jaras, Harada, Bhatt et al., 2025), energy-pool models coupling spiking to local metabolic resources (Bennett, Bhalla and Bhatt, 2017), and pruning strategies targeting energy directly (Chen, Kundu and Beerel, 2024). However, in none of these frameworks does the glutamate/GABA cost asymmetry play an explicit role in shaping learned circuit topology.

A parallel challenge lies in the training of Dale’s-law-compliant excitatory/inhibitory networks (where each neuron maintains a single neurotransmitter phenotype, being either purely excitatory or purely inhibitory). DePasquale et al. (2018) showed that such networks can produce complex temporal dynamics, and Rosenbaum et al. (2017) demonstrated that approximate E/I balance constrains the neural manifold to be approximately linear, while a “semi-balanced” state with selective excess inhibition supports the nonlinear representations needed for nontrivial computation. Structured inhibitory motifs, including vasoactive intestinal peptide (VIP) mediated disinhibition (Pi, Hangya, Kvitsiani, Sanders, Huang, Bhatt and Kepecs, 2013) and interneuron-to-interneuron connectivity (Pfeffer, Xue, He, Huang, Bhatt, Bhatt, Bhatt and Bhatt, 2013), further enrich the computational repertoire. These geometric and dynamical effects are typically derived from stability arguments or functional objectives rather than from explicit resource constraints, leaving open the compelling question of whether metabolic pressure alone can organically organise E/I circuit structure.

In this study, we address these issues by introducing an asymmetric, synapse-type-specific energy cost as a Lagrangian penalty within a surrogate-gradient framework. The key mechanism is that a single penalty term, applied uniformly to all plastic synapses, produces differential gradient pressure on excitatory versus inhibitory pathways purely through the embedded cost asymmetry (*C*_Glut_ > *C*_GABA_) (Fig. 1A). We parametrise the penalty strength by a scalar *λ*_met_ ≥ 0, which we interpret as an effective metabolic pressure. This serves as a systems-level abstraction of global signals (such as adenosine or astrocytic feedback) that dynamically modulate the trade-off between task performance and energetic conservation.

**Figure 1:**
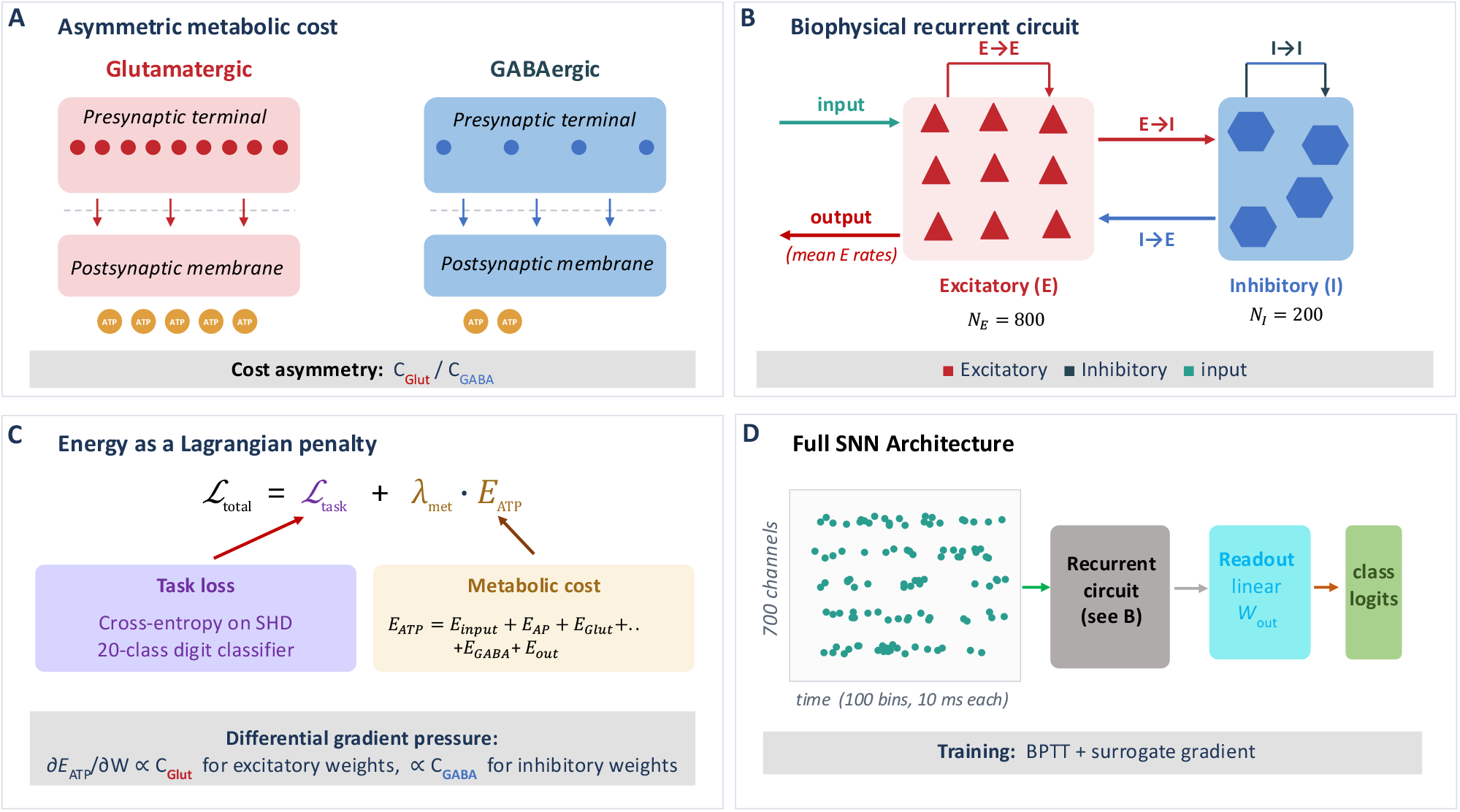
Schematic of the model and training framework. **(A)** Asymmetric metabolic cost model. **(B)** Biophysical recurrent circuit architecture. **(C)** Lagrangian penalty formulation. **(D)** Full SNN architecture and training pipeline.

The implemented SNN comprises a cortical microcircuit with Dale’s-law-compliant excitatory (pyramidal) and inhibitory (fast-spiking Parvalbumin-positive PV^+^) populations, distinct membrane time constants, and four independently plastic recurrent connection classes (*E* → *E, E* → *I, I* → *E*, and *I* → *I*). Crucially, the metabolic penalty extends beyond the recurrent core to both the feedforward input and the linear readout, closing the two routes through which the optimiser could otherwise produce apparent energy savings without genuinely sparsifying the recurrent circuit. The input pathway is penalised to prevent the network from circumventing recurrent computation through unpenalised feedforward drive, and the readout is penalised to prevent the network from exporting energy into an unpenalised decoder (Fig. 1B). We report two main findings:

1. The metabolic penalty functions as a structural regulariser, not merely a cost constraint. The unpenalized network converges into a suboptimal, inhibition-dominated regime with elevated energy and depressed accuracy. Applying mild metabolic pressure moves the network out of this regime, simultaneously improving accuracy and reducing energy several-fold, accompanied by a distinct expansion of representational dimensionality.
2. The asymmetric cost structure induces an emergent pruning order (*E* → *E* most vulnerable, *I* → *E* most resistant) without any pathway-specific prior. The emerging patterns are characterized by excitatory spike costs converging toward the biophysical action-potential floor, while the inhibitory per-spike cost is dominated by the many *I* → *E* connections each surviving interneuron must drive, which makes these neurons individually expensive in dense circuits. These motifs are robust across a two-dimensional sweep of five cost ratios and fifteen structural densities (spanning 75 distinct conditions).

Beyond these principal findings, the constrained optimization formulation itself yields a useful diagnostic toolkit: tracking gradient norms, the gradient balance ratio, and the marginal rate of substitution across the energy/performance (i.e., Pareto) frontier brackets the productive *λ*_met_ band and locates the collapse threshold that bounds it, providing a principled criterion for penalty selection without arbitrary grid searches. Two structural controls reinforce this interpretation: the deliberately low weight ceiling proves non-binding, with synaptic saturation remaining low across all conditions, and the penalised readout is suppressed rather than inflated under pressure, excluding decoder rescaling as an explanation for the observed energy reductions.

Together, these results demonstrate that a single biophysical cost asymmetry is sufficient to organically organise sparse, pathway-specific circuit dynamics in spiking networks. Furthermore, they provide a robust quantitative framework linking metabolic pressure to the emergence of structured excitatory/inhibitory motifs. To our knowledge, this is the first study to use asymmetric metabolic costs as a direct training signal for recurrent spiking networks and to systematically characterise the resulting circuit phenotypes across the Pareto frontier.

## 2. Methods

To computationally formalise the metabolic constraints shaping cortical microcircuits, we translate the biological asymmetry between excitatory and inhibitory energy costs into a differentiable mathematical framework. In biological networks, the metabolic expenditure of synaptic signaling is heavily weighted by the neurotransmitter type, with glutamatergic transmission demanding significantly more ATP than GABAergic transmission. Rather than direclty applying a generic complexity penalty, such as an *L*_1_ or *L*_2_ norm on synaptic weights, we construct a forward model of instantaneous metabolic flux that explicitly scales with these distinct biophysical costs.

This approach allows us to treat energy conservation not merely as an auxiliary constraint, but as a primary driver of circuit topology. By formulating this asymmetric energy model as a Lagrangian relaxation, we can integrate it directly into a surrogate gradient learning framework. Controlling over the global metabolic cost, we consequently apply a control/balance penalty which will naturally exert differential gradient pressures on excitatory versus inhibitory pathways during training. The following subsections detail this mathematical translation, beginning with the formal constrained optimisation problem and the asymmetric energy model that instantiates it, then specifying the recurrent microcircuit, the surrogate-gradient optimisation procedure, and the gradient-balance diagnostics used to characterise the resulting solutions.

### 2.1. Metabolic Penalty as a Lagrangian Relaxation

First, we formalise cortical learning as an optimisation problem subject to energetic constraints. Let ℒ_task_ denote the task error (cross-entropy) and *E*_ATP_ represent the expected metabolic expenditure of the circuit. The constrained problem is defined as:

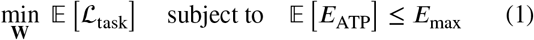

where **W** collects all trainable synaptic weights of the network (the four recurrent connection classes, the feedforward input projection, and the linear readout), expectations are taken over the stimulus distribution and network stochasticity, and *E*_max_ > 0 denotes a fixed upper bound on the per-sample metabolic expenditure (i.e., the total ATP expended during one input presentation), representing the circuit’s effective energy budget. Applying Lagrangian relaxation (Bertsekas, 1999; Boyd and Vandenberghe, 2004) to the constraint in Eq. 1 yields the Lagrangian:

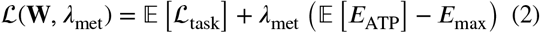

where *λ*_met_ ≥ 0 is the Lagrange multiplier associated with the energy constraint. Because *E*_max_ does not depend on **W**, the term −*λ*_met_ *E*_max_ is a constant that drops out of ∇_**W**_ *L*. We can therefore minimise the reduced penalised objective (Fig. 1C):

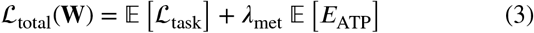

in which the multiplier *λ*_met_ replaces the budget *E*_max_ as the controlled parameter. When *λ*_met_ = 0 the model reduces to standard supervised learning, whereas large *λ*_met_ drives the network towards sparse, low-energy regimes. In a convex loss landscape, Slater’s condition (Boyd and Vandenberghe, 2004) guarantees strong duality, meaning there exists a 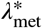 at which the penalised and constrained problems share identical solutions. In our recurrent SNN, by contrast, the task loss is a non-convex function of **W**: the threshold nonlinearity that generates each spike, together with the recurrent dynamics that apply the same weights at every timestep, makes the network output a deeply nested composition of nonlinear maps. Strong duality therefore generally does not hold (Bertsekas, 1999). We therefore treat *λ*_met_ as a penalty coefficient that parametrises a family of objective functions: by sweeping *λ*_met_ we trace a regularisation path whose converged solutions empirically approximate the error/energy Pareto frontier (Boyd and Vandenberghe, 2004).

### 2.2. Asymmetric Synaptic Energy Model

To reflect the biological constraints discussed above, the instantaneous metabolic expenditure at time *t* encompasses the entire circuit, integrating the recurrent core, the feedforward input pathway, and the linear readout:

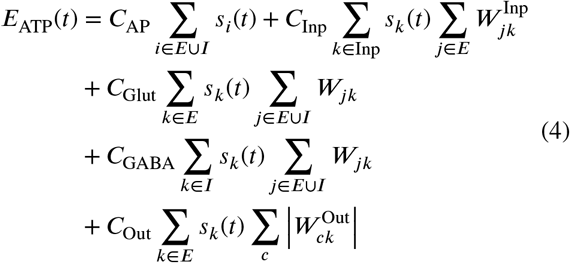

where *s*_i_ (*t*) ∈ {0, 1} represents the binary spike output of neuron *i* at time *t*, 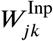 denotes the feedforward inputto-excitatory weight matrix, and 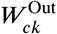 denotes the readout weight from excitatory unit *k* to output class *c*. Action potential costs (*C*_AP_) are exclusively applied to recurrent neurons, as input spikes are generated externally by the cochlear model (Cramer et al., 2022) and fall outside the network’s internal metabolic budget.

In this formulation, the expression (*s*_*k*_ *t* ∑_*j*_ *W*_*jk*_ ) approximates the total synaptic flux activated upon the firing of neuron *k*. Energy is expended only during spike generation and scales proportionally with the aggregate outgoing synaptic conductance. Weight magnitude serves as a proxy for the combined effects of vesicle release probability, receptor density, and postsynaptic conductance. While these are biologically distinct quantities, their associated ATP costs scale approximately with total transmitted synaptic drive (Attwell and Laughlin, 2001). All recurrent and input weights *W*_*ij*_ ≥ 0 are maintained via post-update clamping, with their signs enforced by the Dale’s law convention (detailed in Section 2.3). This ensures that the recurrent and input flux terms are always non-negative and directly interpretable as metabolic costs.

A central feature of this model is the cost asymmetry between neurotransmitter types. Excitatory (glutamatergic) recurrent transmission is metabolically dominant, *C*_Glut_ > *C*_GABA_, with *C*_Glut_ applying to both *E* → *E* and *E* → *I* synapses. All coefficients are expressed in arbitrary units; only their relative ratios affect the dynamics, with numerical values listed in Table 1. Although these are per-synapse coefficients, the total cost incurred when a neuron fires scales with its *fan-out* (the number of postsynaptic targets it connects to): a spike contributes 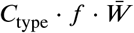 to the energy budget, where *f* is the fan-out, 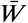 the mean outgoing weight, and *C*_type_ is selected by the presynaptic neurotransmitter. For *f* ≫ 1, this synaptic contribution dominates the somatic action-potential cost *C*_AP_, consistent with experimental estimates of cortical energy expenditure (Attwell and Laughlin, 2001; Sengupta et al., 2010). Under backpropagation, the gradient ∂*E*_ATP_/∂*W*_*ij*_ therefore scales with *C*_Glut_ for excitatory and *C*_GABA_ for inhibitory presynaptic weights, exerting stronger gradient pressure on excitatory pathways.

**Table 1.**
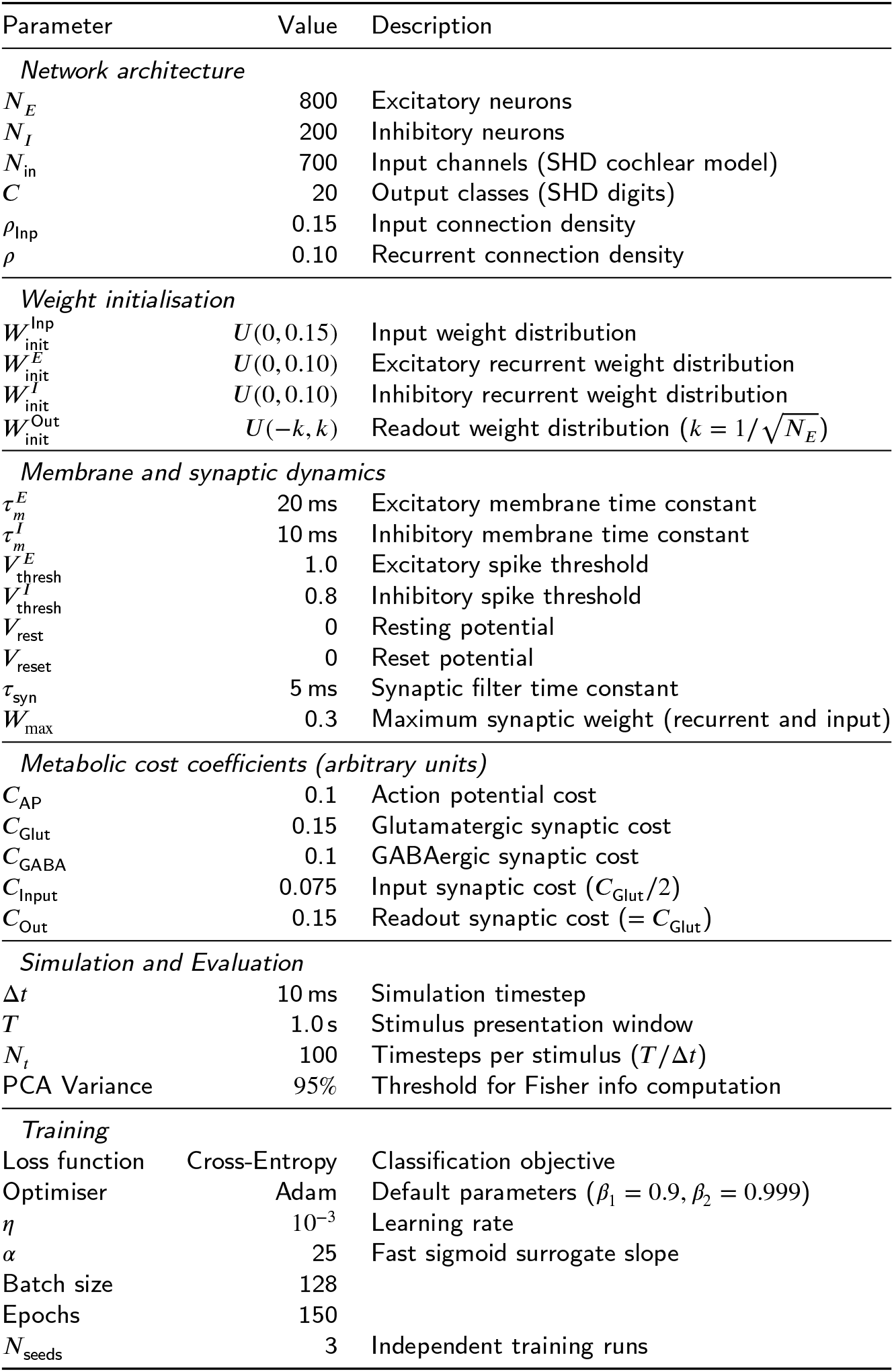
Fixed model parameters.

Feedforward input weights (Inp → *E*) enter *E*_ATP_ at a reduced coefficient *C*_Inp_ = *C*_Glut_/2. Updated cortical energy budgets attribute roughly half of glutamatergic signalling energy to postsynaptic receptor activation and the restoration of postsynaptic ion gradients (Howarth et al., 2012; Attwell and Laughlin, 2001); charging the input pathway at half the recurrent glutamatergic rate therefore retains the postsynaptic share borne by the network while discounting the externally funded presynaptic share. Including the input pathway in the penalty at all is nonetheless an important design choice; without it, the optimiser could collapse recurrent energy to zero while sustaining accuracy entirely through the unpenalised feedforward path.

The final term penalises the linear readout. Each excitatory spike that drives the decoder contributes 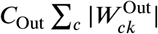,where the absolute value is taken because, unlike the recurrent and input matrices, the readout is signed and is not constrained by Dale’s law. We set *C*_Out_ = *C*_Glut_ (Table 1): the presynaptic units are excitatory, their somatic spikes already incur *C*_AP_, and their projection onto the decoder is glutamatergic, so the readout is charged at the full glutamatergic rate rather than the halved input rate. Penalising the readout closes an offloading route that an input-and-recurrent-only budget would leave open: without it, the optimiser could suppress recurrent synaptic flux while inflating the decoder weights to preserve class separation, exporting the apparent energy saving into an unpenalised pathway. With the readout inside the budget, the decoder is instead driven down under metabolic pressure (Section 3.6), so the reported energy reductions reflect genuine circuit sparsification rather than a rescaling of the readout.

To monitor how the energy is distributed across populations during training, we also track the mean per-spike metabolic cost separately for the excitatory and inhibitory pools:

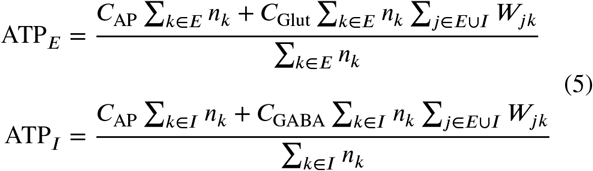

where *n*_*k*_ = ∑_*t*_ *s*_*k*_ (*t*) is the total spike count of neuron . These per-spike measures isolate the recurrent and somatic cost carried by each pool and exclude the readout term, which is reported separately as a budget component. When a population is silent (∑ _*k*_ *n*_*k*_ <0 .5), we default to reporting *C*_AP_ as the floor cost.

### 2.3. Recurrent Cortical Microcircuit Model

We anchor these metabolic penalties within a recurrent network of Leaky Integrate-and-Fire (LIF) neurons featuring distinct excitatory (*E*) and inhibitory (*I*) populations in a 4:1 ratio (*N*_*E*_ = 800, *N*_*I*_ = 200). The two populations possess distinct membrane dynamics to reflect cortical celltype specialisation.

The membrane dynamics of excitatory neurons (pyramidal cells) follow a leaky integrator:

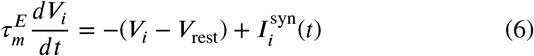

where *V*_*i*_ is the membrane potential of neuron *i, V*_rest_ is the resting potential, 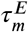 is the excitatory membrane time constant, and 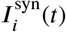 is the total synaptic input at time *t*, expressed in un^*i*^its of membrane potential. Note that the membrane resistance *R*_*m*_ is absorbed into the synaptic drive, so that the physical synaptic current is scaled directly to the voltage it induces. A spike is emitted whenever *V*_*i*_ crosses the excitatory threshold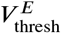, after which is reset to . We set *V*_rest_ = *V*_reset_ = 0; numerical values for all model parameters are listed in Table 1.

In contrast, fast-spiking (PV^+^) inhibitory interneurons possess shorter membrane time constants (indicating higher input conductance) and sit closer to the threshold at rest (Hu, Gan and Jonas, 2014; Gouwens, Sorensen, Berg et al., 2019). We capture this physiological distinction with:

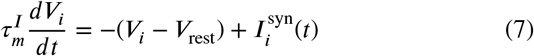

where 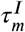 and 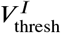 are the inhibitory membrane time constant and^*m*^spike t^t^h^h^r^re^e^s^s^h^hold, both smaller than their excitatory counterparts (Table 1).

For both populations, synaptic input obeys Dale’s law via a sign convention:

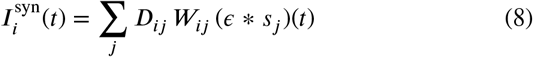

where *D*_*ij*_ = +1 for excitatory and *D*_*ij*_ = −1 for inhibitory presynaptic neurons, *ϵ* represents an exponential synaptic filter with time constant *τ*_syn_, and ∗ denotes convolution. Weights *W*_*ij*_ ≥ 0 are maintained via post-update clamping. The overall recurrent circuit comprises four connection classes (Fig. 1B): *E* → *E, E* → *I, I* → *E*, and *I* → *I*. All four pathways are plastic and subject to the identical gradient rule (detailed below), with no autapses permitted (*W*_*ii*_ = 0 for both *E* → *E* and *I* → *I*). The inclusion of *I* → *I* connections is motivated by the established computational role of interneuron-to-interneuron connectivity in the cortex (Pfeffer et al., 2013; Pi et al., 2013). By allowing them, the optimiser can theoretically route computation through disinhibitory motifs. Because *I* → *I* synapses are governed by *C*_GABA_, they remain metabolically inexpensive. At initialisation, each possible connection within the four recurrent pathways is included independently with probability *ρ*, so that *ρ* fixes the expected fraction of all possible pairwise connections that are realised: the recurrent connection density (Table 1). At *ρ* = 1 a pathway is fully (allto-all) connected, whereas the default *ρ* = 0.10 instantiates roughly one tenth of its possible synapses.

### 2.4. Optimisation via Surrogate Gradients

We optimise the non-differentiable spiking network using BPTT with surrogate gradients. Although BPTT is not biologically local, it reliably finds the weight configurations that minimise the penalised objective for a given *λ*_met_. Whether a biologically local learning rule could converge to the same solution is a separate question outside the scope of this paper. The gradient of Eq. (3) with respect to the weights **W** requires the derivative of the Heaviside step function, which we replace with a continuous surrogate (a fast sigmoid with slope *α*) during the backward pass. The resulting weight update is:

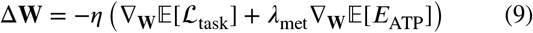

where *η* is the learning rate of the Adam optimiser (Kingma and Ba, 2015). After each update, the recurrent and input weights are clamped to *W*_*ij*_ ∈ [0, *W*_max_] to enforce nonnegativity and a fixed upper bound (values of *α, η*, and *W*_max_ are presented in Table 1). The readout weights *W*^Out^ are exempt from this clamp because the decoder is signed and unbounded, and is constrained only through the metabolic term *C*_Out_. The differential pressure on excitatory versus inhibitory pathways arises automatically from the asymmetric energy model itself, steering the learning trajectory without the need for manual, pathway-specific priors.

The recurrent and input weight ceiling is set relatively low, *W*_max_ = 0.3 (Table 1). At the chosen threshold and synaptic time constant, no single synapse at the ceiling can drive a postsynaptic unit to fire on its own, so spike generation requires the coordinated summation of several inputs and the network cannot satisfy the task by saturating individual connections. Whether this ceiling binds in practice, that is, whether the optimiser parks weights at *W*_max_ rather than eliminating them, is examined directly in Section 3.6.

### 2.5. Gradient Balance at Convergence

To evaluate the equilibrium between task performance and energy conservation, we monitor the gradient balance during training. At an interior stationary point of ℒ_total_, the weight update in Eq. (9) vanishes, so its task and metabolic gradient terms must balance:

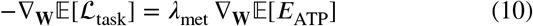

This relationship dictates that the marginal reduction in task error from any weight perturbation is exactly offset by *λ*_met_ times the marginal increase in metabolic cost. While this only strictly holds locally and at an interior optimum in our non-convex, box-constrained setting, we utilize the associated ratio as a diagnostic tool to map regime transitions. Consequently, the gradient balance ratio is defined as:

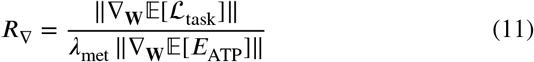

which would converge toward 1.0 at an interior stationary point of the penalised objective. As discussed later in Subsection 3.7, the box constraints [0, *W*_max_] and the large pruned fraction at the lower bound mean that the converged solutions are typically boundary, not interior, optima, so *R*_∇_ need not reach unity even when training has fully converged.

### 2.6. Metric Definitions

We employ several functional metrics to characterise the network’s behaviour under metabolic stress:

The Treves-Rolls lifetime sparsity (Treves and Rolls, 1991; Rolls and Tovee, 1995) measures how selectively neurons respond to specific stimuli:

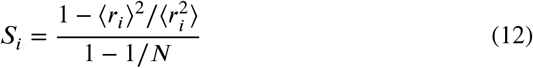

where *r*_*i*_ is the firing-rate response of neuron *i* to a given stimulus, N is the number of stimuli in the evaluation set, and ⟨⋅⟩ denotes the mean over that set. Values approaching zero indicate dense, uniform activation, whereas values near one indicate highly selective, stimulus-specific responses.

To assess the manifold dimensionality of the population response, we calculate the participation ratio (Mazzucato, Fontanini and La Camera, 2016; Gao and Ganguli, 2015):

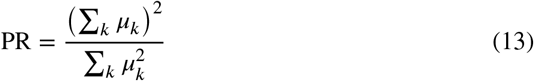

where *µ*_*k*_ are the eigenvalues of the covariance matrix of the mean excitatory firing rates.

The Fisher-discriminability coding efficiency (bits per ATP) (Laughlin et al., 1998; Levy and Baxter, 1996; Harris et al., 2012) is calculated as:

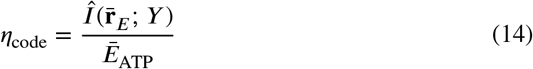

where the mutual-information estimate Î takes the form (Brunel and Nadal, 1998):

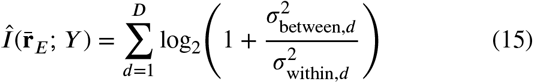

computed after projecting firing rates onto the top *D* principal components retaining 95% of the total variance.

### 2.7. Full Network Architecture and Training

All experiments use the SHD dataset (Cramer et al., 2022), a neuromorphic audio benchmark of spoken digits (*C* = 20 classes) in English and German. The dataset requires genuine temporal computation, as cochlear spike patterns encode temporal structure that cannot be classified from instantaneous snapshots. Spike times are binned into *N*_*t*_ discrete timesteps of width Δ*t* over a stimulus window of duration *T* (Table 1), producing binary spike tensors 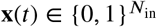 per timestep.

The network architecture, illustrated in Fig. 1D, is organised as a feedforward-recurrent pipeline. An *N*_in_-channel cochlear input layer encodes spoken digits as binary spike tensors, which drive the Dale’s-law-compliant recurrent microcircuit (Section 2.3) of *N*_*E*_ excitatory and *N*_*I*_ inhibitory neurons via a feedforward projection in which each input-to-excitatory connection is present independently with probability *ρ*_input_, the input connection density. The mean excitatory firing rates are then passed through a linear readout 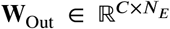 that produces the *C* class logits used to compute the cross-entropy loss. This readout is itself metabolically penalised through *C*_Out_ (Section 2.2). Population sizes, connection densities, and all other architectural parameters are listed in Table 1.

The network is trained end-to-end on cross-entropy loss using BPTT with surrogate gradients (Section 2.4) and the Adam optimiser (Kingma and Ba, 2015); learning rate, batch size, number of epochs, weight bounds, and surrogate slope are listed in Table 1. To control for variance from weight initialisation, all reported metrics are averaged over 3 independent training runs using distinct random seeds, with per-condition values computed as the mean over the final 5 epochs.

## 3. Results

### 3.1. Metabolic Pressure Acts as a Structural Regulariser

To map the trade-off between accuracy and metabolic cost, we trained the SNNs across a logarithmic sweep of *λ*_met_ (16 values in [0, 10^−3^]) at fixed recurrent density *ρ* = 0.10. For each condition, metrics were first averaged over the final 5 epochs of each run, then averaged across *N*_seeds_ independent random seeds (Table 1).

Figure 2A shows validation accuracy as a function of *λ*_met_.The unpenalised network (*λ*_met_ = 0) converges to 70.9%. Although unconstrained surrogate-gradient SNNs routinely achieve 80 to 85% on the SHD benchmark (Cramer et al., 2022; Zenke and Vogels, 2021; Bittar and Garner, 2022; Yin, Corradi and Bohté, 2021; Perez-Nieves, Leung, Dragotti and Goodman, 2021), our baseline’s lower performance is a direct consequence of our biological priors (Dale’s-law compliance alongside biophysically distinct membrane time constants for the excitatory and inhibitory populations), which complicate the optimisation landscape. Introducing mild metabolic pressure improves on this baseline: final-epoch accuracy peaks at 79.5% (*λ*_met_ = 10^−5^), an improvement of 8.6 percentage points. Accuracy then declines steeply at higher penalty strengths as the recurrent circuit collapses.

**Figure 2:**
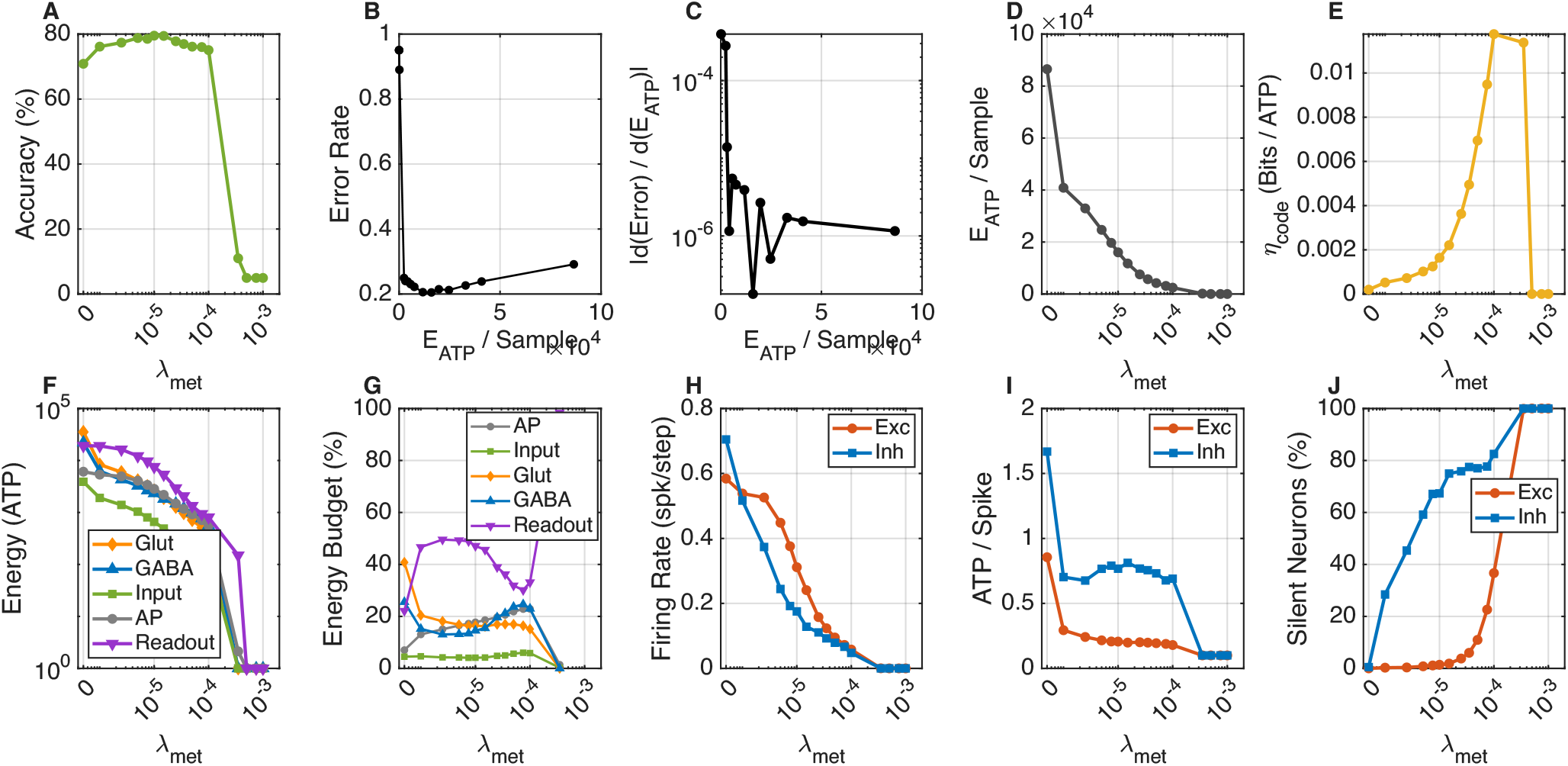
Energy/accuracy trade-off and asymmetric E/I energy allocation across the metabolic sweep. Networks were trained at fixed density *ρ* = 0.10 across 16 values of *λ*_met_ ∈ [0, 10^−3^] for 150 epochs each; all panels report metrics averaged over the final five epochs of each run and then across 3 independent random seeds. **(A)** Validation accuracy. **(B)** Empirical Pareto frontier. **(C)** Marginal rate of substitution on the Pareto frontier. **(D)** Total ATP energy. **(E)** Information efficiency *η*_code_ (bits per ATP). **(F)** Per-component energy decomposition on a logarithmic scale. **(G)** Fractional energy budget. **(H)** Mean firing rates. **(I)** ATP cost per spike. **(J)** Silent neuron fractions.

Inspection of the training trajectory clarifies the suboptimality of the unpenalised baseline: inhibitory firing rates settle at an elevated 0.70 spikes per step, the mean *I* → *E* weight grows steadily over training, and the population code is low-dimensional. Unconstrained learning in this biologically restricted space favours an inhibition-heavy regime that both wastes energy and limits task performance. Mild metabolic pressure is sufficient to move the network out of this basin, so the penalty functions not as a cost constraint but as a structural regulariser that steers the network toward a more productive region of weight space.

Figure 2B plots task error against mean ATP energy across the full sweep, tracing an empirical Pareto frontier with a well-defined knee. The marginal rate of substitution of the frontier, |*∂* (error)/ *∂E*_ATP_|, decays as a power law with increasing energy (Fig. 2C), indicating sharply diminishing returns for metabolic investment beyond the knee.

Figure 2D tracks total ATP expenditure per sample. Without metabolic pressure, the network reaches 86,519 ATP units per sample. Mild pressure cuts this sharply while preserving accuracy: at the peak-accuracy operating point (*λ*_met_ = 10^−5^, 79.5%), total energy falls to 16,019 ATP units per sample, a 5.4× reduction; at the knee used for the two-dimensional sweep below (*λ*_met_ = 5 × 10^−6^, 78.8%), the reduction is 3.5×. Beyond the productive band energy continues to fall by further orders of magnitude, reaching effectively zero at *λ*_met_ = 10^−3^, but only because the circuit silences completely (see Subsection 3.2).

Together, panels A, B, and D of Fig. 2 reveal three distinct operating regimes: a suboptimal baseline at *λ*_met_ = 0 with elevated energy and an inhibition-heavy code; a productive band around the Pareto knee (*λ*_met_ between 10^−6^ and 10^−4^) where accuracy peaks and energy drops several-fold; and a collapsed regime under heavy penalty in which both accuracy and energy fall toward zero as the recurrent circuit dismantles. Figure 2E shows the information-efficiency measure *η*_code_ (bits per ATP). Efficiency peaks within the productive band, at an operating point where accuracy remains 75.1%. The bits-per-ATP optimum therefore nearly coincides with the high-accuracy regime rather than trading accuracy away, so efficiency and task performance are co-optimised across the knee.

Across all five panels, the metabolic penalty consistently acts as a structural regulariser rather than a cost constraint: in the productive band it does not trade accuracy for energy but reorganises the network so that both improve together, until the recurrent circuit collapses under heavy pressure.

### 3.2. Asymmetric Energy Allocation Across Excitatory and Inhibitory Populations

The accuracy and energy gains found in Subsection 3.1 emerge from an asymmetric penalisation of the excitatory/inhibitory spiking activity. Because glutamatergic transmission is more expensive than GABAergic transmission (*C*_Glut_ > *C*_GABA_; Table 1), the optimiser applies different strategies to the excitatory and inhibitory populations. Panels F through J of Figure 2 make this asymmetry visible across five complementary views.

Figure 2F shows the per-component energy budget on a logarithmic scale, now resolved into five terms. At *λ*_met_ = 0, glutamatergic synaptic costs account for 40.8% of total ATP, followed by GABAergic (25.5%), readout (22.1%), action-potential (7.0%), and input (4.5%) costs. As metabolic pressure increases, all five components decline. The glutamatergic, GABAergic and readout terms are eliminated most aggressively; GABA energy drops to zero once all inhibitory neurons are silenced (*λ*_met_ ≥ 3.5 × 10^−4^), and at the strongest penalties every component collapses toward zero as the circuit ceases to spike.

Figure 2G shows the corresponding shift in the fractional composition of the budget, complementing the absolute-magnitude view of Fig. 2F. Because the synaptic components decline faster than the somatic component, the action-potential share rises with *λ*_met_, but only modestly: it climbs from 7.0% at *λ*_met_ = 0 to a maximum of 22.9% near *λ*_met_ = 10^−4^. Rather than converging to an irreducible somatic floor, the circuit silences entirely at the strongest penalties. As the residual absolute budget vanishes instead of becoming spike-dominated, the fractional budget briefly becomes dominated by the readout (reaching 100%), which reflects the cost of the final few excitatory spikes driving the decoder just before complete silencing. This is a direct consequence of the low weight ceiling: with *W*_max_ = 0.3, surviving units cannot sustain firing once their synaptic drive is pruned, and neurons fall silent rather than continuing to spike through stripped connections (Subsection 3.6).

Figure 2H shows the mean firing rates of the two populations. Both rates decline with increasing *λ*_met_, but the inhibitory population starts from an elevated baseline (0.70 spikes per step) and is compressed far more steeply, consistent with the preferential suppression of inhibition described below.

Figure 2I traces the per-spike cost explicitly. The excitatory cost declines from 0.86 at *λ*_met_ = 0 to 0.10 at the strongest penalties, converging exactly onto the somatic floor *C*_AP_ (Table 1). This indicates that the optimiser has pruned essentially all excitatory synaptic flux, leaving each surviving excitatory unit operating at the minimum metabolic cost per spike. The inhibitory per-spike cost is highest at the baseline (1.67 at *λ*_met_ = 0), reflecting the elevated inhibitory rate combined with strong *I* → *E* fan-out weights, and it falls monotonically as inhibition is suppressed. A fan-out-driven rise in inhibitory per-spike cost does occur, but along the structural-density axis of the two-dimensional sweep (Subsection 3.5) rather than along *λ*_met_.

Figure 2J quantifies the silencing phenomenon, defining a silent neuron as one that emits zero spikes during the stimulus presentation window. The fraction of silent inhibitory neurons rises steeply with *λ*_met_, reaching 100% by *λ*_met_ = 3.5 × 10^−4^. Excitatory silencing proceeds much more gradually, remaining low through the productive band before accelerating to 100% at *λ*_met_ = 10^−3^. The near-complete elimination of inhibition occurs while accuracy remains high (Fig. 2A), demonstrating that the optimiser discovers that the SHD task can be sustained by a predominantly excitatory microcircuit, until the strongest penalties silence the excitatory pool as well.

Together, panels F to J of Figure 2 reveal an asymmetric allocation of metabolic cost across the two populations: the optimiser does not compress activity uniformly, but selectively eliminates the inhibitory population while preserving and sparsifying the excitatory one toward the biophysical floor, a partition that follows directly from the embedded cost asymmetry *C*_Glut_ > *C*_GABA_.

### 3.3. Pathway-Specific Pruning and Representational Geometry

Figure 3A tracks the aggregate excitatory-to-inhibitory synaptic ratio, Σ*W*_*E*_/Σ*W*_*I*_ . The ratio is non-monotonic. It starts at 1.33 in the unpenalised network and first eases toward parity, reaching a minimum of *∼*1.12 near *λ*_met_ = 10^−5^ as glutamatergic drive is pruned preferentially. It then rises to a peak of *∼*1.42 near *λ*_met_ = 7.5 × 10^−5^, as the onset of inhibitory silencing reduces the aggregate inhibitory weight faster than the excitatory, before collapsing toward zero at the strongest penalties as the surviving weights are eliminated.

**Figure 3:**
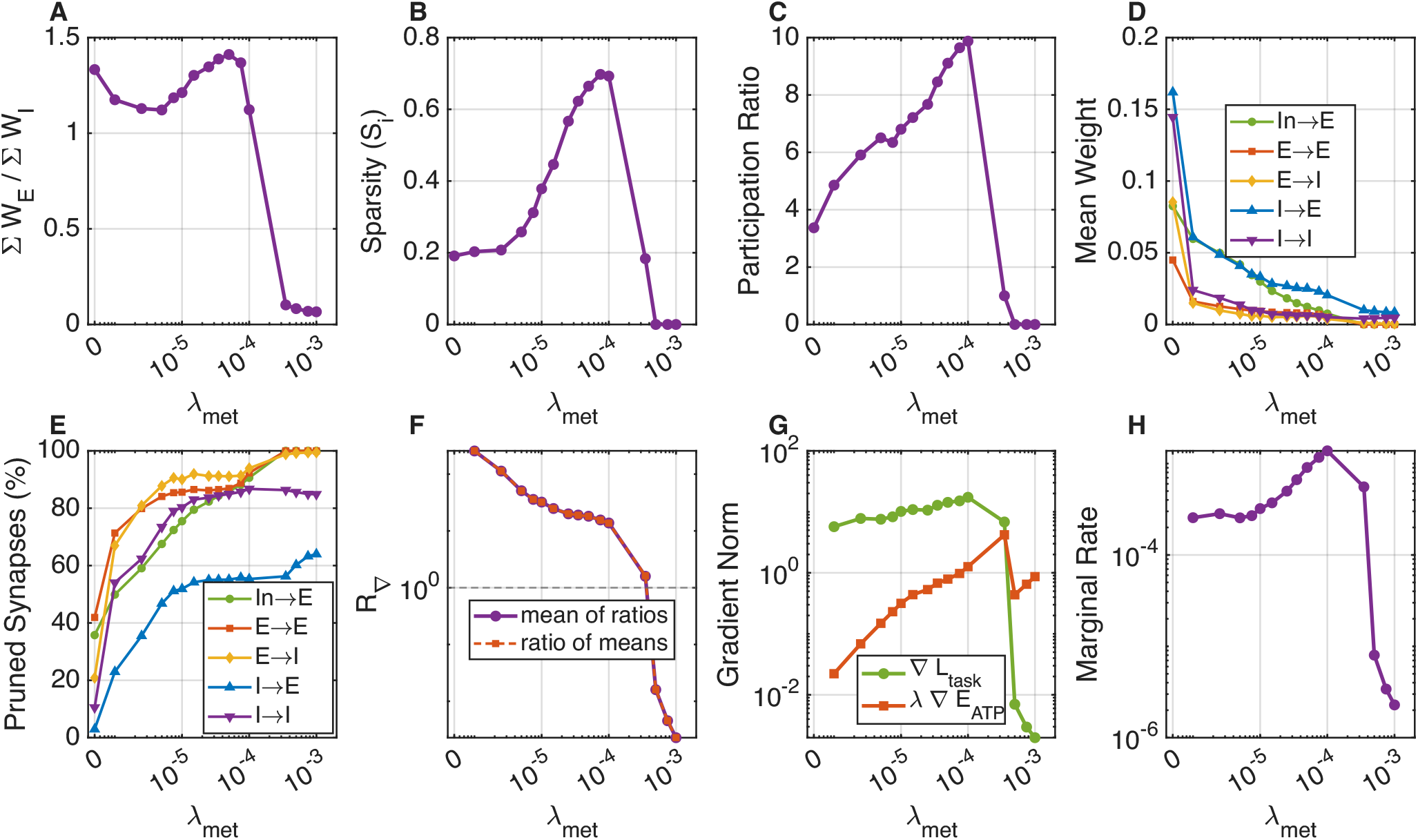
Pathway-specific synaptic reorganisation, representational geometry, and Lagrangian diagnostics. Same *λ*_met_ sweep as Figure 2. All panels report metrics averaged over the final five epochs of each run and then across 3 independent random seeds. **(A)** E/I weight ratio. **(B)** Lifetime sparsity. **(C)** Participation ratio. **(D)** Mean synaptic weight by connection type. **(E)** Pruning fractions. **(F)** Gradient balance ratio *R*_∇_ on a logarithmic axis; the dashed line marks unity. Circles show the mean across seeds of the per-seed ratio; squares show the ratio of the seed-averaged gradient norms of panel G. The two aggregations differ by at most 2.2%, confirming that the comparison between panels F and G is unaffected by averaging order. **(G)** Gradient norms. **(H)** Marginal rate of substitution.

Figure 3B shows the Treves-Rolls lifetime sparsity. It rises from near zero in the unpenalised network to a peak of 0.70 at *λ*_met_ = 7.5 × 10^−5^, indicating progressively sparser population codes as redundant neurons are silenced, with a slight relaxation under extreme starvation as the few surviving neurons must each carry more representational load.

Figure 3C shows the participation ratio, an estimate of the effective dimensionality of the population response manifold. The profile is also non-monotonic. At *λ*_met_ = 0 the participation ratio is low (*∼*3.4), reflecting the inhibition-heavy baseline code. As metabolic pressure increases, the participation ratio rises to a peak of *∼*9.9 near *λ*_met_ = 10^−4^, indicating that moderate metabolic pressure physically expands the effective dimensionality of neural representations by relieving the inhibition-dominated regime and distributing activity across a wider subspace. Under stronger penalties the participation ratio declines again as too few neurons remain active to support a high-dimensional code. The expansion of representational dimensionality coincides with the productive band, suggesting that the metabolic regulariser improves task performance in large part by enlarging the computational manifold available to the readout layer.

Figure 3D shows the mean weight of each pathway across the sweep. At baseline the inhibitory pathways carry the largest mean weights, having grown during unpenalised training; input and excitatory weights start lower and the input weights decline steeply under the dedicated input penalty. Among the recurrent pathways, *I* → *E* connections preferentially retain synaptic drive, maintaining the largest mean weight through intermediate *λ*_met_, consistent with the preferential preservation of inhibitory feedback onto excitatory targets even as overall density falls.

Figure 3E shows the corresponding pruning fractions, which follow a clear neurotransmitter-ordered hierarchy. Glutamatergic and input pathways are the most vulnerable: *E* → *E* connections are pruned most aggressively (71.3% by *λ*_met_ = 10^−6^, 100% at 10^−3^), with *E* → *I* (67.0% to 99.4%) and input (49.9% to 100%) close behind. The GABAergic pathways resist longest, and among all pathways *I* → *E* is preserved the most: it is only 23.0% pruned at *λ*_met_ = 10^−6^ and 64.0% at 10^−3^, while *I* → *I* is pruned 54.0% to 84.9% over the same range. The persistence of the inhibitory pathways is itself a window into the limits of the metabolic penalty: because GABA transmission is cheaper per synapse these connections pass through the cost filter most easily, but above *λ*_met_ = 3.5×10^−4^ all inhibitory neurons are silenced, leaving the surviving inhibitory weights (chiefly *I* → *E*) functionally inert. We refer to these as “zombie synapses”, since the penalty on weight magnitude alone is insufficient to drive them to zero once their presynaptic neurons have ceased firing.

Together, panels A to E of Fig. 3 reveal a structural reorganisation that is both pathway-specific and representationally productive: differential pruning eliminates excitatory recurrent and input drive most aggressively while preserving inhibitory feedback to excitatory targets, and the resulting circuit operates on an expanded population manifold that underwrites the high-accuracy regime identified in Subsection 3.1.

### 3.4. Lagrangian Diagnostics

To confirm that the structural reorganisations described above emerge from a controlled trade-off between the task and metabolic objectives, we tracked the gradient norms of both terms throughout training. As established in Section 2.5, the gradient balance ratio *R*_∇_ (Eq. 11) would approach unity at an interior stationary point of the penalised objective, and its magnitude reports which of the two objectives dominates the update at a given penalty strength.

Figure 3F tracks *R*_∇_ across the sweep on a logarithmic axis, with the dashed line marking unity. Because *R*_∇_ is a ratio, its average across seeds is not uniquely defined: the mean of the per-seed ratios and the ratio of the seed-averaged norms differ in general, and can differ substantially when the distribution across seeds is heavy-tailed. We report the mean of the per-seed ratios, each seed first averaged over its final five epochs. The two aggregations agree closely here, differing by at most 2.2% across the sweep and by less than 0.6% throughout the productive band, so nothing that follows depends on the choice of aggregation. Upon activation of the metabolic penalty the ratio falls across more than five orders of magnitude, from *∼*256 at *λ*_met_ = 10^−6^ to *∼*18 at 5 × 10^−5^ and *∼*0.002 at 10^−3^, where the circuit has silenced. The descent is monotonic at every step, with no reversals, consistent with a single continuously deformed family of solutions rather than jumps between qualitatively different optima. It is not uniform, however: the ratio remains far above unity throughout the productive band and falls below it only once the circuit begins to collapse.

Figure 3G decomposes the same quantity into its numerator and denominator. Throughout the productive band the task gradient exceeds the metabolic term by one to two and a half orders of magnitude, and *R*_∇_ crosses unity only between *λ*_met_ = 3.5 × 10^−4^ and 5 × 10^−4^, by which point the inhibitory population is already fully silenced. This is expected rather than anomalous: as noted in Section 2.5 and revisited in Subsection 3.7, the box constraints [0, *W*_max_] and the large pruned fraction at the lower bound place the converged solutions on the constraint boundary, so the interior stationarity condition that would drive *R*_∇_ → 1 does not apply.

The diagnostic value of *R*_∇_ therefore lies in the shape of the curve rather than its absolute level. Across the decade from *λ*_met_ = 10^−5^ to 10^−4^ the ratio flattens into a plateau, falling only from ∼32 to ∼14, whereas the single step from 3.5 × 10^−4^ to 5 × 10^−4^, a factor of 1.4 in *λ*_met_, drops it by two orders of magnitude. The plateau identifies the band over which the regularisation path is stable, and the sharp break beneath it locates the collapse threshold.

Figures 2C and 3H show the marginal rate of substitution |∂ (error) / ∂*E*_ATP_ |, plotted against total energy and against *λ*_met_ respectively. As a function of energy (Fig. 2C), the sensitivity decays as a power law: above the Pareto knee, each additional unit of energy buys progressively less accuracy. As a function of *λ*_met_ (Fig. 3H), the same quantity rises through the productive band, peaking near the knee (*λ*_met_ ≈ 10^−4^), before falling steeply in the collapse regime, where the silenced circuit becomes insensitive to further penalty. The monotonic increase up to the knee confirms that each additional unit of metabolic penalty extracts a progressively larger accuracy cost, and supports the convexity of the empirical Pareto frontier in the regime of interest.

Together, the four panels (Fig. 2C and Fig. 3F,G,H) support the constrained-optimisation interpretation of the regularisation path. Non-convexity and the box constraints preclude formal claims of global optimality or of *R*_∇_ → 1. Nonetheless, the monotonic descent of *R*_∇_ and the marginal rate of substitution provide a mathematically grounded criterion for selecting *λ*_met_: the plateau in *R*_∇_, together with the peak in the marginal rate of substitution, brackets the productive band and locates the collapse threshold that bounds it, reducing the need for arbitrary grid searches.

### 3.5 Robustness Across Cost Ratios and Structural Densities

The *λ*_met_ sweep established the regularising effect of metabolic pressure at a single density and cost ratio. To test whether the resulting circuit motifs generalise, we conducted a two-dimensional sweep crossing five cost-asymmetry ratios (*C*_Glut_/*C*_GABA_ ∈ {1.0, 1.25, 1.5, 1.75, 2.0}) with fifteen initial recurrent densities (*ρ* ∈ [0.05, 1.0]), all evaluated at fixed *λ*_met_ = 5 × 10^−6^ (chosen near the Pareto knee identified in Subsection 3.1). This yields 75 distinct conditions, shown in Fig. 4 (functional metrics) and Fig. 5 (pathway-specific connectivity).

**Figure 4:**
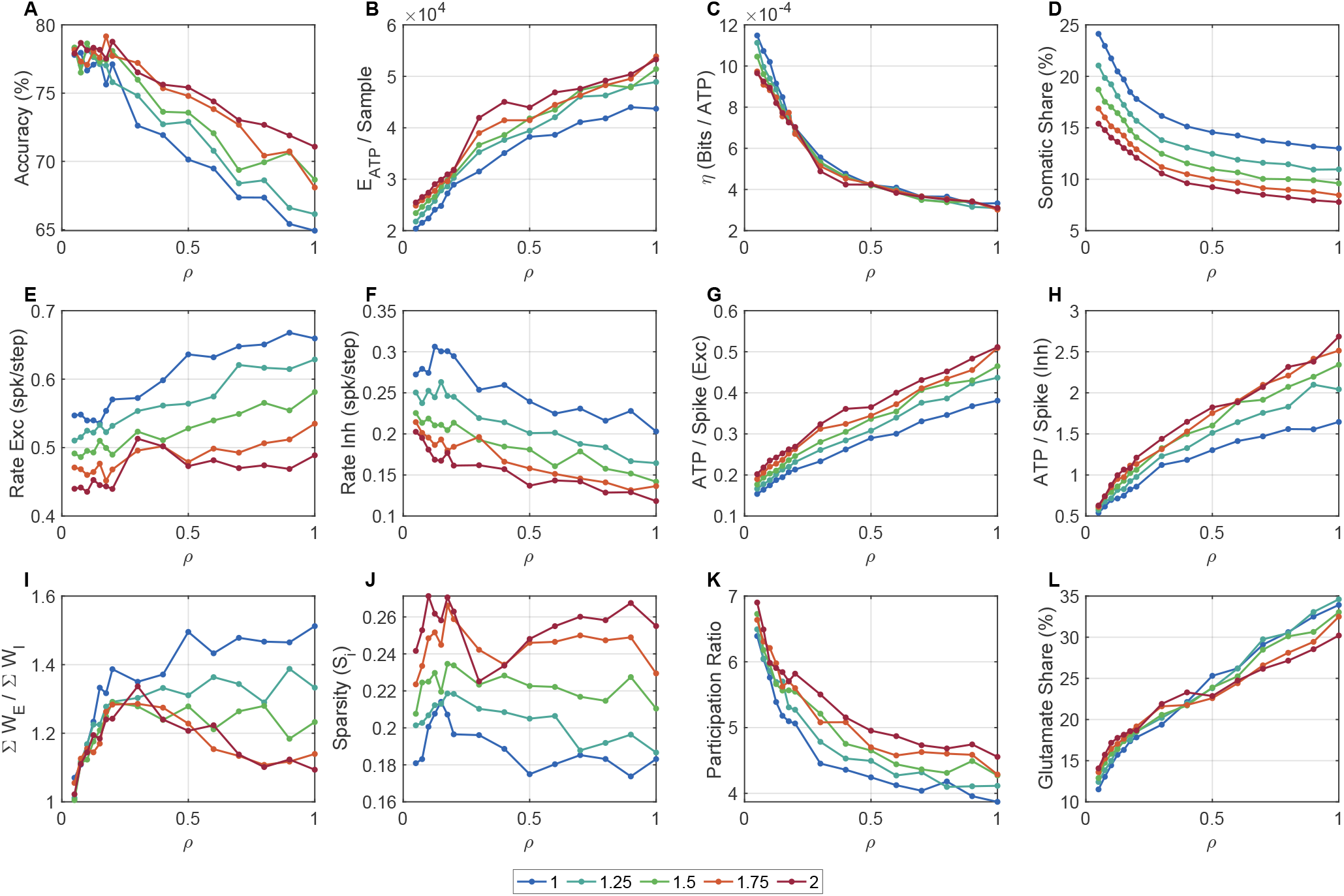
Performance, energy budget, and representational geometry across the cost-ratio × density sweep. Networks were trained at fixed *λ*_met_ = 5 × 10^−6^ across 5 cost ratios and 15 densities. All panels report metrics averaged over the final five epochs of each run and then across 3 independent random seeds. **(A)** Validation accuracy. **(B)** Total ATP energy. **(C)** Information efficiency *η*_code_ (bits per ATP). **(D)** Somatic (action-potential) share of total energy. **(E, F)** Excitatory and inhibitory mean firing rates. **(G)** Excitatory ATP per spike. **(H)** Inhibitory ATP per spike. **(I)** E/I weight ratio. **(J)** Lifetime sparsity. **(K)** Participation ratio. **(L)** Glutamatergic share of total energy.

**Figure 5:**
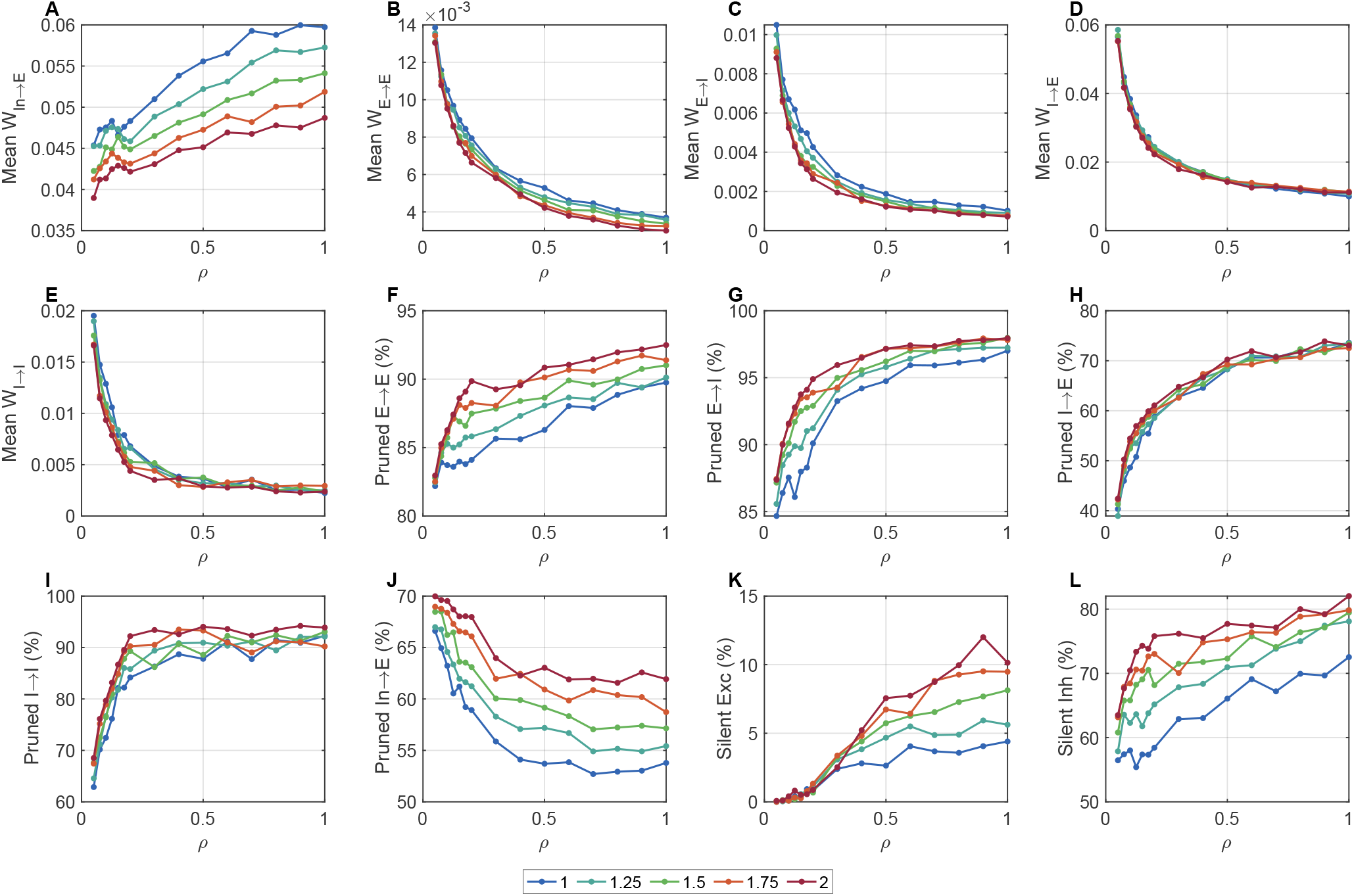
Pathway-specific synaptic reorganisation across the cost-ratio × density sweep. Same conditions as Figure 4. All panels report metrics averaged over the final five epochs of each run and then across 3 independent random seeds. **(A to E)** Mean synaptic weights by pathway. **(F, G)** Glutamatergic pruning fractions (*E* → *E* and *E* → *I*). **(H, I)** GABAergic pruning fractions (*I* → *E* and *I* → *I*). **(J)** Input pruning fraction. **(K)** Silent excitatory neuron fraction. **(L)** Silent inhibitory neuron fraction.

Figure 4A reports validation accuracy across all 75 conditions. Accuracy remains remarkably robust, ranging from 64.9% to 79.2% (mean 74.3% ± 3.9%). The five cost-ratio curves overlap almost entirely, confirming that accuracy is largely insensitive to the degree of cost asymmetry at this *λ*_met_. Density exerts a modest but systematic effect: sparse networks (*ρ* ≤ 0.15) average 77.7% accuracy, while dense networks (*ρ* ≥ 0.60) average 69.8%, reflecting how the metabolic penalty disproportionately compresses the computational repertoire of densely connected circuits. Figure 4C shows the complementary information efficiency *η*_code_ (bits per ATP), which declines gradually with density as the larger metabolic footprint of denser circuits diminishes their per-spike informational return.

Figure 4B tracks total post-optimisation energy. Although the initial ATP footprint scales linearly with *ρ* (synapse count is proportional to density), after 150 epochs of optimisation the energy gap between the sparsest and densest networks compresses to a factor of only ∼2.1×. At cost ratio 2.0, for example, energy rises from ∼25,478 ATP at *ρ* = 0.05 to ∼53,279 at *ρ* = 1.0, only a 2.1× increase despite 20× more initial synaptic hardware. Figure 4D,L reveal where this sublinearity comes from: the metabolic budget shifts progressively from somatic toward synaptic with increasing density. At cost ratio 2.0, the somatic (action-potential) share of total energy falls from 15.4% at *ρ* = 0.05 to 7.8% at *ρ* = 1.0 (panel D), while the glutamatergic share grows from 14.1% to 30.2% over the same range (panel L), reflecting the higher fan-out per spike in denser networks. Both shifts hold across all five cost ratios.

Figure 4E,F show that the asymmetric E/I energy allocation described in Subsection 3.2 generalizes across all conditions. Excitatory firing rates remain in a narrow band across the density range, while inhibitory rates decline from ∼0.20 at *ρ* = 0.05 to ∼0.12 at *ρ* = 1.0. Figure 4G,H show the corresponding per-spike costs. The excitatory cost rises modestly from ∼0.20 to ∼0.51 across the density range, while the inhibitory cost surges from ∼0.63 at *ρ* = 0.05 to ∼2.68 at *ρ* = 1.0. This density-driven rise, absent along the *λ*_met_ axis (Fig. 2I), directly reflects the increasing *I* → *E* fan-out concentrated on the few remaining active interneurons in denser circuits.

Figure 4I shows that the silencing-driven aggregate asymmetry also generalises: the Σ*W*_*E*_/Σ*W*_*I*_ ratio rises modestly with density (from ∼1.0 at *ρ* = 0.05 to ∼1.1 at *ρ* = 1.0), paralleled by a rise in inhibitory silencing from ∼63.5% to ∼82.0% across the same range (Fig. 5L). Figure 4J,K characterise the representational geometry: lifetime sparsity remains largely invariant to density (panel J), while the participation ratio declines systematically from ∼6.9 at *ρ* = 0.05 to ∼4.6 at *ρ* = 1.0 (panel K), indicating that the dimensionality of the population code contracts as the underlying connectivity densifies.

Figure 5 confirms that the pathway-specific pruning hierarchy established in Subsection 3.3 also generalises across the entire two-dimensional sweep. Panels A through E display the mean synaptic weights for each pathway, demonstrating a consistent retention of *I* → *E* synaptic drive relative to excitatory recurrent connections across all densities. The corresponding pruning fractions (panels F to J) reproduce the neurotransmitter-ordered hierarchy: glutamatergic pathways are consistently devastated, with *E* → *E* pruning bounded between 82.2% and 92.5% and *E* → *I* between 84.7% and 98.0% regardless of the initial density. In contrast, the GABAergic pathways show a much broader dynamic range and are the most resistant. *I* → *E* pruning varies from 38.9% in sparse networks to 73.9% in the densest, and *I* → *I* from 62.9% to 94.2%, confirming their relative metabolic shielding under sparse-to-moderate conditions. Input synapses (panel J) are pruned at a stable rate of 52.7% to 70.0%.

Finally, panels K and L explicitly contrast the population silencing phenomena. Excitatory silencing (panel K) remains low across all 75 evaluated conditions, reaching at most 12.0% in the most aggressive corner of the sweep. Inhibitory silencing (panel L), however, scales strongly with density, rising from 63.5% to 82.0%. Notably, while the macroscopic functional metrics in Figure 4 were largely insensitive to the specific cost ratio, panel L reveals that higher cost ratios accelerate this inhibitory silencing phenomenon, pushing it further toward completion as density increases. This confirms that the embedded per-synapse asymmetry directly drives the selective elimination of the inhibitory population.

Together, these results establish the asymmetric E/I energy allocation and the pathway-specific pruning hierarchy as generic consequences of the metabolic regulariser: they hold across all 75 conditions, with density acting as the dominant axis of quantitative variation and cost asymmetry as a secondary modulator.

### 3.6. Weight-Ceiling Occupancy and Readout Cost

Two design choices in the energy model invite a direct control: the low weight ceiling *W*_max_ = 0.3, which could in principle drive the optimiser to park weights at the boundary rather than eliminate them, and the readout penalty *C*_out_, which is intended to close the decoder-offloading route described in Section 2.2. Figure 6 examines both directly across the *λ*_met_ sweep.

**Figure 6:**
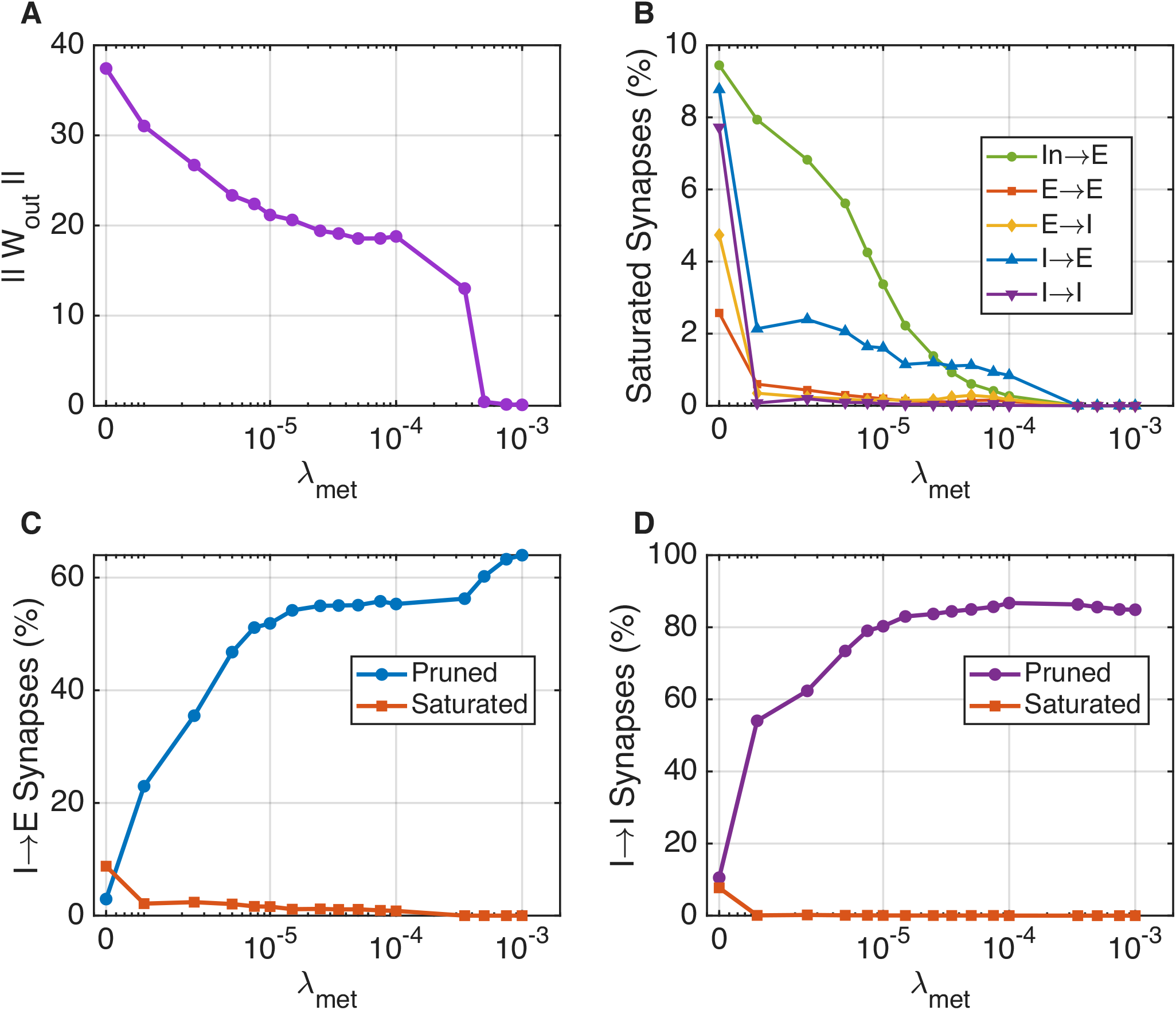
Weight-ceiling occupancy and readout cost across the metabolic sweep. Same *λ*_met_ sweep as Figure 2. All panels report metrics averaged over the final five epochs of each run and then across 3 independent random seeds. **(A)** Readout norm ‖ **W**_out_‖. **(B)** Fraction of surviving synapses at the weight ceiling (≥ 0.95 *W*_max_) by pathway. **(C)** *I* → *E* pruned versus saturated fraction. **(D)** *I* → *I* pruned versus saturated fraction.

Figure 6A shows the readout norm ‖**W**_out_ ‖as a function of *λ*_met_. Far from inflating to absorb the energy removed from the recurrent core, the readout norm *falls* under metabolic pressure, from ∼37.4 in the unpenalised network to ∼0.1 at the strongest penalties. The penalty therefore suppresses the decoder rather than the optimiser exploiting it as an unpenalised reservoir, confirming that the energy reductions in Subsection 3.1 reflect genuine circuit sparsification and not a rescaling of the readout.

Figure 6B shows the fraction of surviving synapses sitting at the weight ceiling (*W*_*ij*_ ≥ 0.95 *W*_max_) for each pathway. Saturation is low throughout: it never exceeds 9.4% in any pathway across the entire sweep, with the input pathway (In → *E*) highest at 9.4%, *I* → *E* at 8.8%, *I* → *I* at 7.7%, and the recurrent excitatory pathways below 5%. The ceiling is therefore non-binding: the optimiser eliminates connections rather than parking them at *W*_max_, so the pruning hierarchy of Subsection 3.3 reflects genuine elimination of synaptic flux rather than an artefact of the chosen bound.

Figures 6C and 6D make this explicit for the two most resistant pathways, contrasting the pruned fraction against the saturated fraction for *I* → *E* and *I* → *I*. In both pathways the pruned fraction rises steeply with *λ*_met_ while the saturated fraction remains low across the entire sweep. The resistance of *I* → *E* is therefore a genuine preservation of inhibitory feedback, not weights pinned at the ceiling, and the rising inhibitory per-spike cost reported along the density axis (see Subsection 3.5) reflects fan-out concentration on surviving synapses rather than ceiling saturation.

The same weight-ceiling control holds across the twodimensional sweep. Figure 7 shows that saturation remains low at every condition: the input pathway is highest at 5.0% to 10.0% across densities, while the recurrent pathways stay near zero (*E* → *E* and *E* → *I* below 0.5%, *I* → *E* below 4.6%, *I* → *I* below 0.3%). The readout norm likewise stays bounded across the grid (panel F), ranging only from ∼21.9 to ∼32.0. The ceiling is therefore non-binding and the decoder bounded across the full two-dimensional sweep, so the pruning hierarchy and the energy reductions reflect genuine elimination of synaptic flux rather than ceiling saturation or decoder offloading.

**Figure 7:**
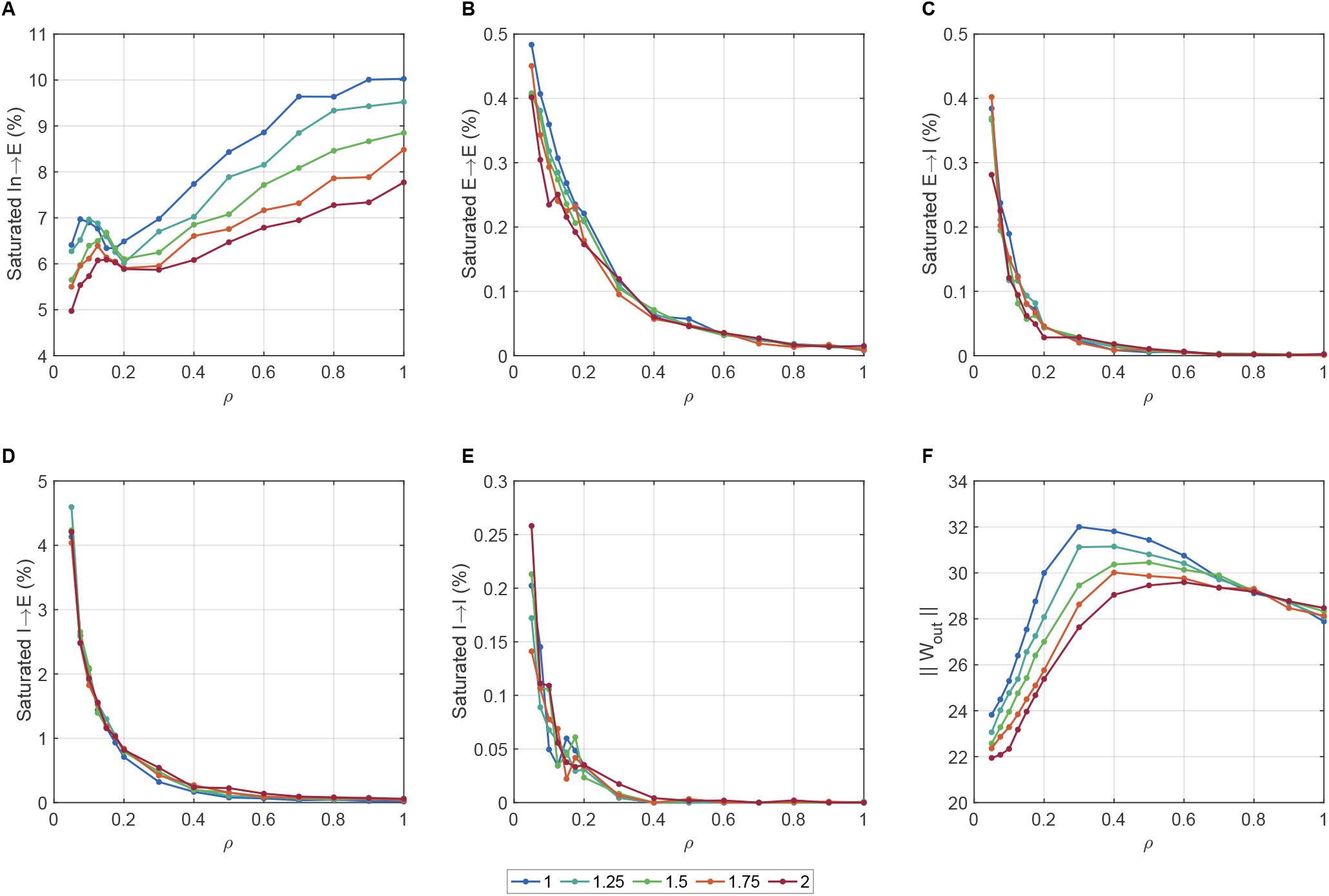
Weight-ceiling occupancy and readout cost across the cost-ratio × density sweep. Same conditions as Figure 4. All panels report metrics averaged over the final five epochs of each run and then across 3 independent random seeds. **(A to E)** Fraction of surviving synapses at the weight ceiling (≥ 0.95 *W*_max_) by pathway. **(F)** Readout norm ‖**W**_out_‖.

### 3.7. Training Convergence and the Epoch Budget

To verify that the structural quantities reported above reflect converged solutions rather than transient states, we tracked full training trajectories over the 150-epoch budget for five representative penalty strengths (Fig. 8).

**Figure 8:**
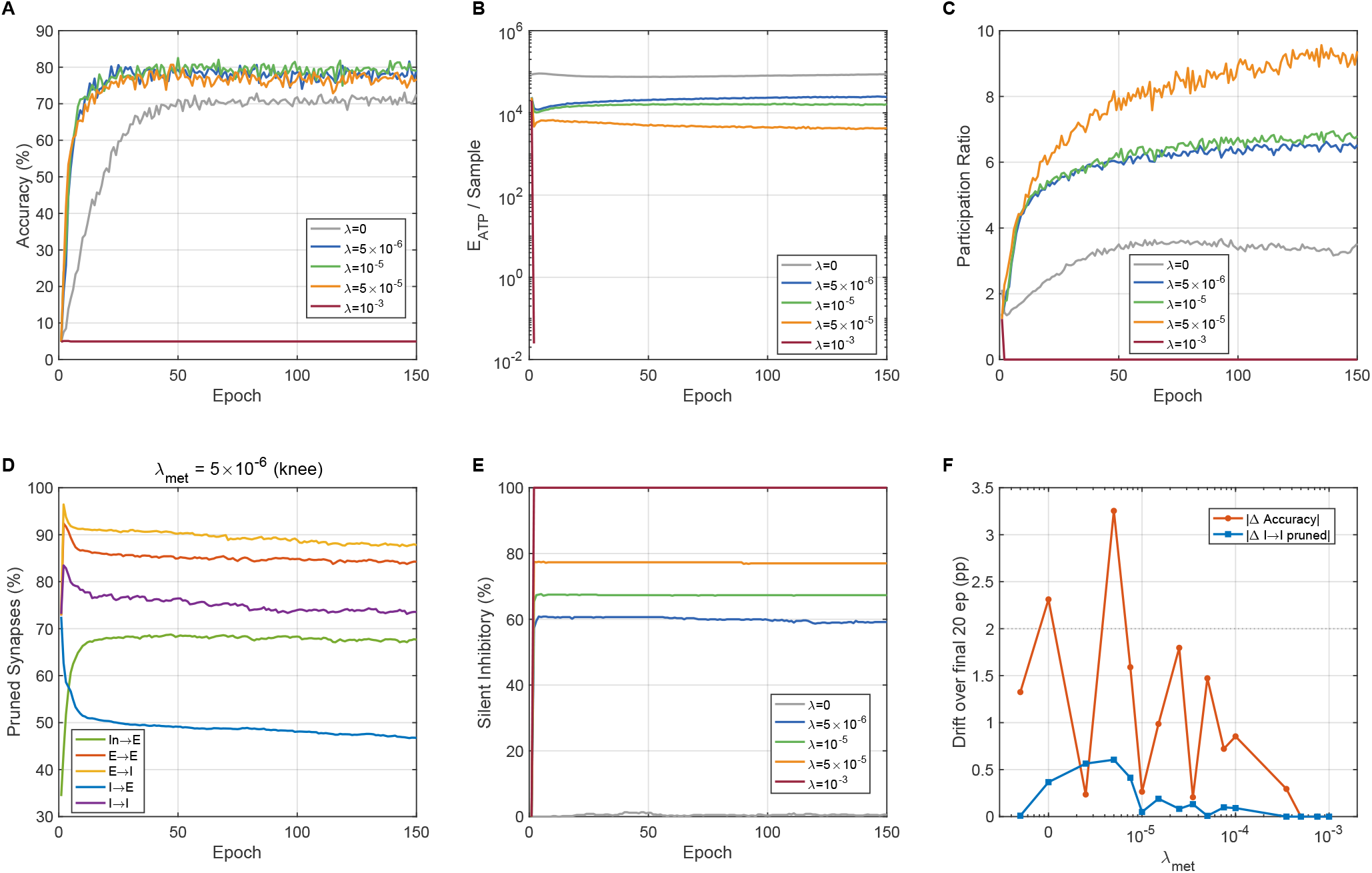
Training convergence over the 150-epoch budget. Trajectories are shown for five representative penalty strengths. All panels report values averaged across 3 independent random seeds. **(A)** Validation accuracy. **(B)** Energy per sample (log scale).**(C)** Participation ratio. **(D)** Per-pathway pruning fractions at the Pareto knee (*λ*_met_ = 5 × 10^−6^). **(E)** Silent inhibitory neuron fraction. **(F)** Drift in accuracy and *I* → *I* pruning over the final 20 epochs.

Figure 8A shows validation accuracy against epoch. In the productive band, accuracy rises quickly and then plateaus well within the budget, with the unpenalised baseline and the heavily penalised runs bracketing the productive curves. Figure 8B shows that energy settles even faster: total ATP per sample drops sharply over the first few tens of epochs and is essentially flat thereafter, except for the unpenalised run, whose energy continues to grow over training, confirming the runaway tendency of the unconstrained baseline. Figure 8C shows that the participation ratio stabilises on a similar timescale.

Figure 8D tracks the per-pathway pruning fractions at the knee penalty (*λ*_met_ = 5 × 10^−6^). The pruning hierarchy is established early and is stable for the remainder of training, indicating that the connectivity reported in Subsection 3.3 is a converged property of the solution rather than a snapshot of ongoing pruning. Figure 8E shows the corresponding inhibitory silencing trajectories, which likewise plateau within the budget.

Figure 8F summarises convergence directly, plotting the drift in accuracy and in *I* → *I* pruning over the final 20 epochs as a function of *λ*_met_. Across the productive band the drift is small, below roughly one percentage point, so the metrics reported here (averaged over the final five epochs) are representative of the converged state. The only appreciable late-training drift occurs at the unpenalised baseline, whose energy is still growing, and in the collapsing tail, where the circuit is progressively silencing; neither regime is the operating point of interest. We note that the gradient balance ratio *R*_∇_ (Subsection 2.5) is not expected to reach unity even at this converged state, because the box constraints [0, *W*_max_] and the large pruned fraction at the lower bound place the solution on the constraint boundary, where the stationarity condition that motivates *R*_∇_ → 1 does not apply.

## 4. Discussion

We set out to ask whether the measured cost asymmetry between glutamatergic and GABAergic transmission is sufficient, on its own, to organise biologically recognisable circuit structure in a trained spiking network. The answer is yes: a single scalar penalty, applied uniformly to all plastic synapses, produces pathway-specific pruning hierarchies, population-level silencing phenomena, and representational geometry changes that are robust across a wide range of cost ratios and structural densities.

### 4.1. Metabolic Pressure as a Regulariser, Not Just a Constraint

Perhaps the most notable finding from our experiments is that metabolic pressure actively improves task performance rather than merely constraining it. The unpenalised network occupies a suboptimal, inhibition-heavy basin: inhibitory firing rates are elevated, *I* → *E* weights grow over training, and the population code is low-dimensional. Introducing metabolic pressure moves the network out of this basin, raising accuracy while simultaneously reducing energy severalfold. This is not the behaviour expected of a simple cost constraint, which would typically trade accuracy for efficiency. Instead, the metabolic penalty acts as a structural regulariser that reshapes the loss landscape, steering the optimiser toward a more productive region of weight space.

Mechanistically, this regularising effect has a natural interpretation in terms of excitatory/inhibitory dynamics. In the unpenalised network, strong *I* → *E* drive suppresses excitatory neurons into a narrow, low-dimensional activity manifold, so the readout has little discriminative signal to work with. Metabolic pressure preferentially suppresses this overactive inhibitory population because each inhibitory spike is expensive in total ATP at baseline, given the elevated inhibitory rate and strong fan-out, despite GABA being cheaper per synapse. As inhibition is reduced, excitatory activity becomes more heterogeneous and stimulus-specific, the participation ratio rises, and the readout gains access to a higher-dimensional code.

Broadly, this phenomenon resonates with existing theoretical work on E/I balance and neural coding. Denève and Machens (2016) argued that tight E/I balance produces efficient codes by cancelling predictable activity, leaving only informative residuals. Similarly, Rosenbaum et al. (2017) showed that approximate balance constrains spiking networks to approximately linear manifolds, while selective excess inhibition (“semi-balance”) enables the nonlinear representations needed for complex tasks. Our results add a new dimension to this picture: metabolic pressure does not preserve a perfectly balanced state, but progressively reduces inhibition, allowing the network to discover that the benchmark task can be sustained by a predominantly excitatory circuit. This aligns with recent observations in neuromorphic engineering where generic, synapse-agnostic regularisation often over-constrains network dynamics; our biologically grounded asymmetry provides a principled alternative that organically expands representational capacity.

### 4.2. Pathway-Specific Pruning and the Zombie Synapse Problem

Building on these population-level dynamics, the emergent pruning hierarchy follows directly from the embedded cost asymmetry but carries nontrivial consequences. Glutamatergic connections experience stronger gradient pressure and are therefore pruned first, with *E* → *E* the most vulnerable pathway. GABAergic pathways are pruned more slowly, and among all four recurrent classes the inhibitory feedback pathway *I* → *E* is preserved the longest.

Within this hierarchy, the persistence of inhibitory synapses under extreme pressure deserves particular attention. A substantial fraction of inhibitory synapses, chiefly *I* → *E*, survive even under heavy metabolic pressure, long after all inhibitory neurons have been completely silenced. These connections are functionally inert. Because their presynaptic neurons no longer fire, they contribute nothing to either the energy cost or the task computation, yet the metabolic penalty on their weight magnitude alone is insufficient to drive them to zero. We termed these “zombie synapses”. Their existence reveals a specific limitation of the current energy model. In conventional deep learning, magnitude-based pruning algorithms routinely sweep away such weights. However, in our biologically inspired framework, a synapse whose presynaptic neuron is silent incurs zero energy cost, so the penalty has no remaining gradient with which to remove it. Crucially, two controls confirm that this persistence is genuine rather than an artefact of the weight bound. First, the weight-ceiling occupancy analysis (see Subsection 3.6) shows that surviving synapses are not parked at *W*_max_: saturation never exceeds 9.4% in any pathway, so *W*_max_ is non-binding and the resistant pathways reflect genuine preservation. Second, the readout norm falls rather than inflates under pressure, excluding the possibility that the apparent recurrent sparsification is offset by decoder rescaling. Resolving the zombie synapse problem would therefore require incorporating the metabolic cost of synaptic maintenance (Harris et al., 2012), which is independent of electrical activity, rather than tightening the weight bound.

On the other end of the spectrum, the convergence of excitatory spike costs exactly onto the biophysical action-potential floor is a striking result. It indicates that the optimiser prunes essentially all excitatory synaptic flux, leaving each surviving excitatory neuron operating at the minimum possible metabolic cost per spike. This is highly consistent with the bits-per-ATP optimisation principle proposed by Harris et al. (2012) and Laughlin et al. (1998): under metabolic pressure, the network converges toward a regime where information is transmitted as cheaply as possible. The inhibitory per-spike cost does not rise along the *λ*_met_ axis; the fan-out-driven rise in inhibitory cost is instead a property of structural density, discussed next.

### 4.3. Density and Cost Ratio: Robustness of the Circuit Motifs

Crucially, the two-dimensional sweep confirms that these circuit-level phenomena are not artefacts of a particular parameterisation. The pruning hierarchy, inhibitory silencing phenomenon, and representational geometry changes are qualitatively preserved across all conditions, with accuracy remaining remarkably robust. This robustness confirms that the metabolic regulariser defines a broad functional basin in weight space that remains accessible from diverse structural starting points.

Across these parameters, structural density (*ρ*) emerges as the dominant axis of variation. Denser networks incur higher total energy, exhibit more complete inhibitory silencing, and support lower-dimensional representations. The sublinear scaling of post-optimisation energy with density demonstrates that the optimiser aggressively compensates for structural redundancy, primarily by silencing inhibitory neurons and pruning excitatory synaptic flux. This compensatory mechanism is functionally analogous to homeostatic plasticity in biological networks, where circuits adjust their excitability to maintain stable function despite large variations in underlying synaptic drive or structural connectivity. It is also along this density axis that the inhibitory per-spike cost rises sharply, from ∼0.63 to ∼2.68 ATP per spike as density increases at fixed cost ratio. This rise reflects *I* → *E* fan-out concentrated on the few surviving interneurons in denser circuits, and stands in contrast to the *λ*_met_ sweep at fixed density, where the inhibitory per-spike cost is highest at baseline and falls under pressure. The same fan-out mechanism therefore produces opposite trends along the two axes, depending on whether the number of surviving inhibitory targets is being driven by density or by penalty strength.

The cost asymmetry ratio plays a secondary role, modulating the intensity of differential pruning without changing its qualitative character. Even without a built-in asymmetry, the pruning hierarchy persists because the fundamental difference in population firing rates and fan-out structure still produces differential total costs per spike between excitatory and inhibitory neurons. However, intentionally increasing the asymmetry widens the gap between excitatory and inhibitory silencing, confirming that the per-synapse cost asymmetry amplifies a structural pattern already present due to networklevel dynamics.

### 4.4. Lagrangian Diagnostics and Practical Implications

Moving beyond structural observations, the gradient balance diagnostics provide a practical tool for selecting the metabolic penalty strength. The upper part of the productive band appears as a plateau in *R*_∇_, spanning roughly a decade over which the ratio changes by only a factor of two, and is bounded at its upper end in *λ*_met_ by an abrupt two-orderof-magnitude break that marks the onset of circuit collapse. We stress that *R*_∇_ is a diagnostic rather than a measure of stationarity: the task gradient dominates the metabolic term throughout the productive band, and because the converged solutions are box-constrained and heavily pruned they sit on the constraint boundary, where the condition motivating *R*_∇_ → 1 does not hold.

The monotonic increase of the marginal rate of subtitution across the sweep supports the convexity of the empirical Pareto frontier in the regime of interest, confirming that each additional unit of metabolic penalty extracts a progressively larger accuracy cost. This property argues against the observed regime transitions being mere artefacts of the optimiser becoming trapped in qualitatively different local optima, and supports the interpretation that a single, smoothly varying family of solutions connects the unpenalised and high-penalty regimes. For practical applications in neuromorphic engineering, such as mapping algorithms to event-driven hardware like BrainScaleS-2 (Cramer et al., 2022), this indicates that the energy/accuracy trade-off is both predictable and tuneable: a system designer can select a point on the Pareto frontier according to their power budget, with reasonable confidence that small changes in the metabolic penalty produce proportionate, rather than catastrophic, changes in performance, provided one remains within the productive band and away from the collapse threshold.

### 4.5. Limitations

Our conclusions are subject to several limitations. First, the steep accuracy collapse beyond the productive band indicates that the circuit reorganization dynamics are sensitive to penalty strength near the collapse threshold; the network transitions from a high-accuracy sparse regime to complete silencing over a relatively narrow range of *λ*_met_. Learning rate schedules, gradient clipping, or adaptive penalty strategies may smooth this transition and are worth exploring.

Second, the energy model utilised here is a first-order approximation. The synaptic flux term captures the dominant scaling of ATP cost with synaptic drive but omits several biologically relevant factors. These include presynaptic vesicle depletion and recycling dynamics, astrocytic spatial constraints, and the highly nonlinear relationship between firing rate and oxidative ATP demand. Incorporating these factors could alter the details of the pruning hierarchy and an explicit activity-independent cost for maintaining synaptic connections would resolve the zombie synapse problem that magnitude penalties alone cannot.

Third, we deploy BPTT with surrogate gradients as a normative optimisation tool, not as a direct model of biological learning. Consequently, the circuit motifs we report primarily characterise the solutions that a globally optimal learner would find under metabolic constraints. Whether biologically local learning rules, such as e-prop (Bellec et al., 2020) or three-factor plasticity models, reliably converge to similar solutions remains an open and vital question. If they converge to the same pruning hierarchy and inhibitory silencing phenomenon, this would strengthen the normative interpretation of our results.

### 4.6. Future Directions

Investigating the connection between metabolic pressure and representational dimensionality on more complex tasks and architectures represents a promising frontier. The observation that moderate metabolic pressure expands the computational manifold while improving accuracy echoes theories of neural efficiency, which propose that cortical circuits operate near a metabolically constrained capacity limit (Harris et al., 2012). Scaling this microcircuit to larger networks with multiple interneuron subtypes (PV, SST, VIP), or training it on tasks that explicitly require disinhibitory control (Pi et al., 2013), would test whether the progressive elimination of inhibition observed here is a general consequence of metabolic pressure, or a phenomenon specific to the task and circuit complexity studied in this work.

## Acknowledgements

This work was partially supported by grant RGPIN-2022-03042 from Natural Sciences and Engineering Council of Canada.

## Data availability

The Spiking Heidelberg Digits (SHD) dataset analyzed in this study is a publicly available benchmark and can be freely accessed via its official repository (Cramer et al., 2022). The Python scripts used to generate the results and replicate the study is available from the corresponding author upon request.

## CRediT authorship contribution statement

**Roberto C. Sotero:** Conceptualization, Methodology, Software, Writing. **Jose M. Sanchez-Bornot:** Methodology, Writing.

**Roberto C. Sotero** received the B.Sc. degree in Nuclear Physics from the Higher Institute for Nuclear Sciences and Technologies (ISCTN), Cuba, in 2003, and the Ph.D. degree in Physics from Havana University, Cuba, in 2009. From 2010 to 2014, he was a postdoctoral fellow at the Montreal Neurological Institute, McGill University, Canada. Since 2014, he has been a faculty member with the Department of Radiology, University of Calgary, Canada, where he is currently an Associate Professor and Principal Investigator who leads the Computational Neurophysics Lab. His research interests include computational neuroscience, neuroimaging, machine learning, and complex networks.

**Jose M. Sanchez-Bornot** received the B.Sc. degree in Computer Science from the University of Havana, Cuba, in 2003, and the Ph.D. degree in Applied Mathematics from the Cuban Neuroscience Centre in 2011. He is currently a Research Software Engineer at Ulster University, U.K. He has more than 20 years of experience in computational neuroscience, neuroimaging, and scientific computing. His research interests include EEG/MEG source localization, functional and effective connectivity, signal processing, machine learning, high-dimensional neuroimaging analysis, and computational modelling of brain activity. He has worked on methods for large-scale brain data analysis, including connectivity estimation, statespace modelling, and machine-learning pipelines for neurological and neurodegenerative disorders

